# The basal ganglia transform visual identity into behavioral relevance

**DOI:** 10.64898/2026.04.07.716881

**Authors:** Julie MJ Fabre, Matteo Carandini, Andrew J Peters, Kenneth D Harris

**Author notes:** Co-senior authors (Matteo Carandini, Andrew J Peters, Kenneth D Harris).

## Abstract

The basal ganglia are thought to be essential for transforming sensory signals into appropriate actions, yet it remains unclear how this transformation unfolds across their nuclei. We recorded neuronal activity across the striatum, globus pallidus external (GPe), and substantia nigra pars reticulata (SNr) in mice that were either naive or trained in visuomotor associations. In naive mice, visual stimuli drove responses in all three nuclei. These responses were feature-selective in the striatum but less so in GPe and SNr. Task training potentiated visual responses across the circuit, particularly for stimuli associated with movement. Training affected nuclei differently: while the striatum continued to distinguish between stimuli associated with the same behavioral action, GPe and SNr did not, responding similarly to all “Go” cues while distinguishing them from “No-Go” cues. These results demonstrate that the basal ganglia progressively transform high-dimensional sensory representations into low-dimensional signals of behavioral relevance.

## Introduction

The basal ganglia play an important role in sensory-motor associations. The striatum, the basal ganglia’s primary input nucleus, shows sensory responses in naive subjects^1–5^, which become potentiated when stimuli are associated with behavioral responses^6–10^. These striatal representations play a causal role in the associated behaviors^11–14^, presumably reflecting their propagation through other basal ganglia nuclei to either cortical^15,16^ or subcortical^17–22^ targets involved in producing the actions. Striatal outputs converge topographically on substantia nigra pars reticulata (SNr), either directly, or indirectly via the globus pallidus external (GPe)^23–26^. These direct and indirect streams have been classically understood as implementing “Go” and “No-Go” pathways^27–29^.

To understand the basal ganglia’s role in sensory-motor association, however, we need to understand if responses to sensory stimuli propagate across their nuclei, and how these responses change when stimuli become behaviorally relevant. The SNr contains roughly 100 times fewer neurons than the striatum^30,31^, leading to the hypothesis that information becomes condensed as it flows into and across the basal ganglia, for example discarding stimulus details that are not behaviorally important^32,33^. Consistent with this possibility, neurons in the striatum can show stimulus-specific responses^34–37^, while neurons in the SNr have been reported to reflect learned stimulus value^38,39^.

To address these questions, we recorded visual responses across the striatum, GPe, and SNr in mice which were either naive or had learned a visuomotor association task that dissociated stimulus identity and behavioral contingency. In naive mice, neurons were driven primarily by large azimuth-spanning stimuli, with stimulus selectivity decreasing from striatum to GPe and SNr. Task training substantially increased visual responsiveness, in a manner that differed between nuclei: after training, striatal cells better distinguished stimulus identity, while GPe and SNr distinguished only the behavioral association (Go vs. No-Go), showing stronger responses to Go stimuli. Thus, learning increases visual sensory responses in the basal ganglia, with information about stimulus identity transformed into information about behavioral relevance across the basal ganglia nuclei.

## Results

### Visual responses along an anatomical basal ganglia pathway in naive mice

To investigate basal ganglia responses to visual stimuli in naive mice, we presented stimuli on three screens positioned to the left, front, and right of head-fixed mice, spanning approximately 270° of visual angle in azimuth and 70° in elevation (**Fig. 1a**). We recorded from the striatum, GPe, and SNr with Neuropixels probes (**Supp. Fig. 1**). Units were identified using an automated curation algorithm^40^ (**Supp. Fig. 2**) to ensure unbiased selection of high-quality single units. To isolate visual responses from motor-related activity, we excluded trials with detectable wheel movements from analysis (**Supp. Fig. S3**). We first investigated which levels of the basal ganglia contain neurons driven by visual stimuli in naive mice by analyzing responses to static gratings presented on all three screens simultaneously (**Fig. 1a**), which drove responses in naive mice across all three regions. These stimuli generally increased the firing rate of striatal neurons (**Fig. 1b,c**, top). In GPe, they increased the firing rate of some neurons and decreased that of other neurons (**Fig. 1b,c**, middle). Similar results were seen in SNr, where suppression was even more common (**Fig. 1b,c**, bottom). These responses were also observed for smaller gratings and full-screen natural images (**Supp. Fig. 4**).

**Figure 1:**
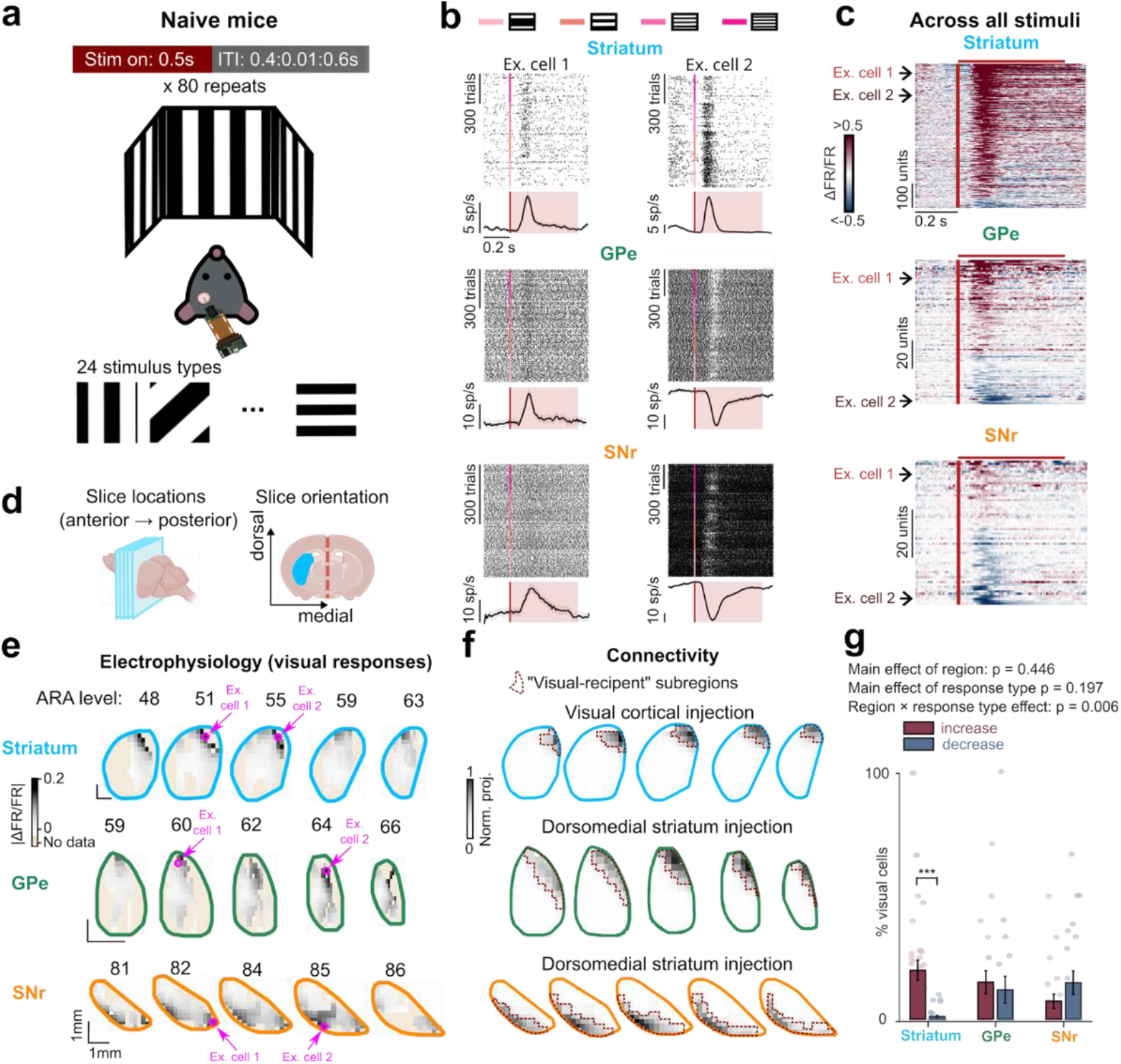
Visual responses along an anatomical basal ganglia pathway in naive mice. **(a)** Head-fixed mice viewed visual stimuli (0.5 s duration, 0.4-0.6 s inter-trial interval) on three screens spanning 270° of visual angle in azimuth. Twenty-four stimulus types were shown including gratings of varying spatial frequency and orientation, natural images, and fractals (Fig. 2c, Supp. Fig. 5). N = 56 sessions in 10 mice (striatum), 32 sessions in 10 mice (GPe), 31 sessions in 8 mice (SNr). **(b)** Raster plots and PSTHs (mean firing rate ± s.e.) of two example neurons from each region (striatum, GPe, SNr, top to bottom). Within each region, rasters are shown above and PSTHs below. Trials are sorted by spatial frequency (all orientations combined), from darker to lighter pink: 1/7.5, 1/10, 1/15, and 1/30 degrees/cycle. Pink lines indicate stimulus onset time. Shaded boxes indicate stimulus duration. **(c)** Pseudocolor PSTHs of all visually responsive neurons’ responses to gratings (averaged over grating spatial frequency and orientation), with neurons sorted vertically by response magnitude (average z-scored response 50 to 200 ms after stimulus onset on even trials, odd trials are plotted here). Arrows: example cells from (b). Red vertical bars indicate stimulus onset. Red horizontal bars indicate stimulus duration. Visually responsive neurons were identified by permutation tests between baseline and post-stimulus onset firing rate (p < 0.01, 999 permutations). Of n = 1737 neurons in striatum, 737 significantly increased in rate, 70 significantly decreased; of n = 522 neurons in GPe, 87 significantly increased and 40 significantly decreased; of n = 254 neurons in SNr, 28 significantly increased, 35 significantly decreased. **(d)** Schematic illustrating the slice locations shown in (e) and (f). Left: slices are arranged anterior to posterior from left to right. Right: example coronal slice at the level of the striatum; cyan fill indicates an example recorded (e) or anatomically defined visual-recipient (f) striatal slice; red dashed line indicates the midline. **(e)** Anatomical distribution of visually responsive neurons across the basal ganglia, shown in coronal sections ordered from anterior (left) to posterior (right). Section positions are indicated by Allen Reference Atlas (ARA) level number, which indexes the standard set of coronal reference plates (lower numbers = more anterior). Pixel darkness indicates the magnitude of absolute firing rate change relative to baseline (|ΔFR/FR|), averaged across all neurons recorded at each location. Magenta circles and arrows mark locations of example cells shown in (b). **(f)** Anatomical distribution of visual cortical projections to the basal ganglia, data from the Allen Connectivity Atlas^41^. Top row: mean axonal projection density in the striatum, averaged across 265 experiments with AAV tracer injections in visual cortical areas. Bottom two rows: mean axonal projection density in GPe and SNr, averaged across 13 and 8 experiments, respectively, with tracer injections in dorsomedial striatum. Red dashed outlines indicate “visual-recipient” domains defined by thresholding the projection matrices from visual cortices for striatum and from “visual-recipient” striatum for GPe and SNr **(g)** Fraction of visually responsive neurons in visual subregions of each nucleus (p < 0.01, permutation test between baseline and post-stimulus firing rate). Each point shows fraction of cells from one recording from the visual subregion of one nucleus that significantly increase (red) or decrease (blue) rate in response to the stimulus. Two-way ANOVA showed no main effect of region (p = 0.45) or response type (p = 0.20) but did show region × response type interaction (p = 0.0057). Post-hoc t-tests showed more neurons excited than inhibited in striatum (p = 0.0009) but not GPe or SNr. Striatum: n = 341 neurons, 30 sessions, 7 mice in the visual subregion. GPe: n = 193 neurons, 19 sessions, 9 mice. SNr: n = 155 neurons, 26 sessions, 7 mice. Error bars: s.e.

Visually responsive neurons, however, were found only in specific portions of those regions: the dorsomedial portion of striatum, the dorsal portion of GPe, and the ventral portion of SNr (**Fig. 1d,e**), which we hypothesized correspond to an anatomical visual pathway that processes input from visual cortex. We evaluated this hypothesis by querying the Allen Mouse Brain Connectivity Atlas^41^ for all projections from visual cortical areas to striatum, and from visual-recipient striatal subregions to GPe and SNr (**Supplementary Table 1**). As hypothesized, the subregions of striatum containing visually responsive units (**Fig. 1e**, top row) closely matched the subregions receiving visual cortical input (**Fig. 1f**, top row). In turn, the subregions of GPe and SNr containing visually responsive units closely matched the subregions receiving input from visual dorsomedial striatum (**Fig. 1e,f**, middle and bottom rows).

Within the visually responsive portions of each nucleus, similar fractions of neurons showed visual responses across all three nuclei (**Fig. 1g**). However, the fraction of positively vs. negatively responding neurons differed across regions (p = 0.0057, region × response type sign ANOVA interaction). Visual responses in striatum were mostly increases in firing rate, with significantly more increases than decreases (p = 0.0009, post-hoc t-test). GPe exhibited increases or decreases in firing rate with roughly equal proportions. SNr showed a trend toward more decreases, though this was not significant.

### Visual selectivity in naive mice decreases along the basal ganglia nuclei

Having established that neurons in a visual pathway through the basal ganglia respond to full-field gratings, we next investigated how selective neurons in this pathway are for visual stimulus features. To do so, we analyzed responses to the battery of visual stimuli: twenty-four stimulus types including gratings with varying spatial frequencies and orientations, natural images, and fractals (**Fig. 2c**, **Supp. Fig. 5**).

**Figure 2:**
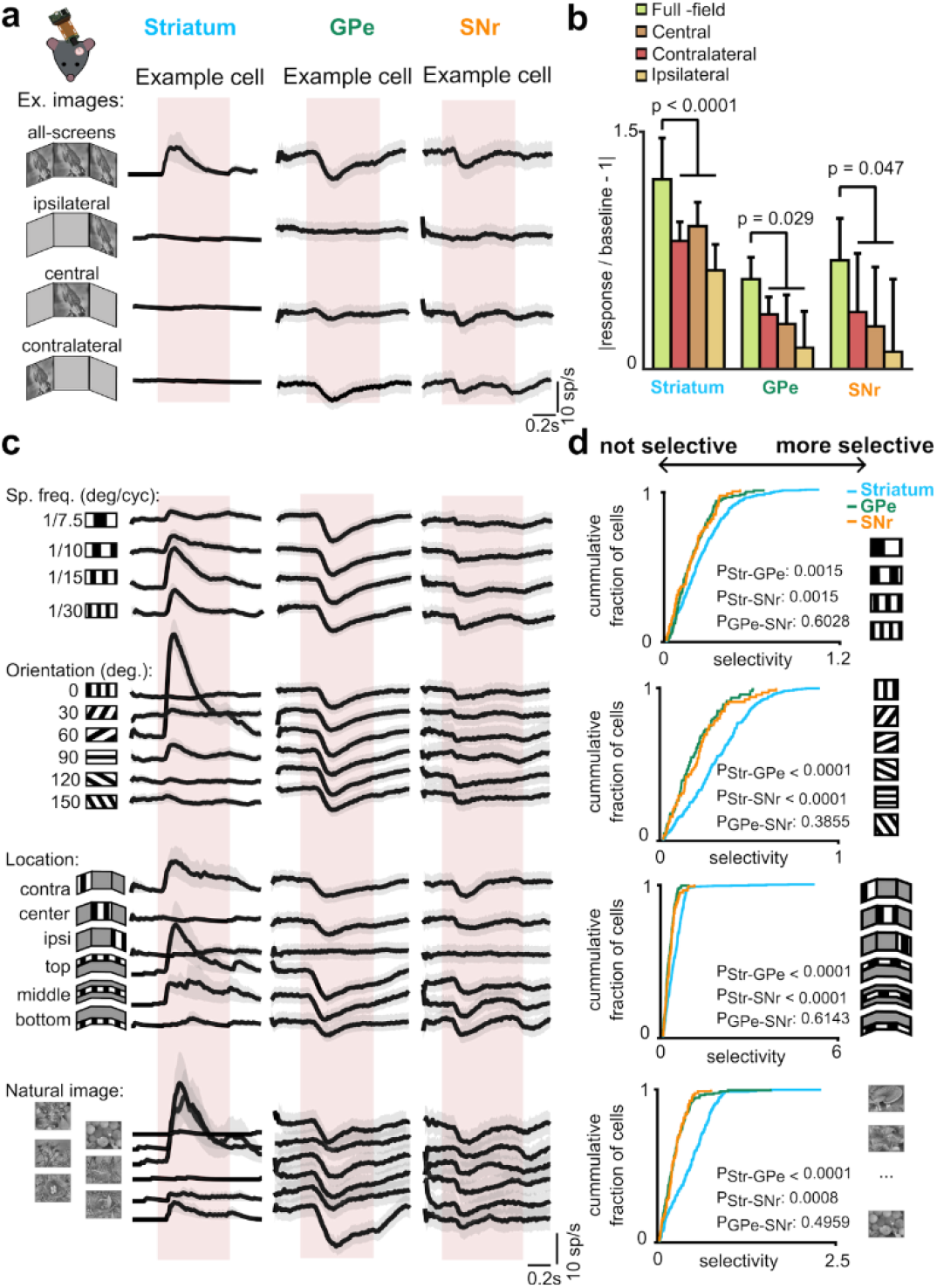
Visual selectivity in naive mice decreases along the basal ganglia nuclei. **(a)** Responses (mean ± s.e. across trials) of example neurons from striatum, GPe, and SNr to one example natural image, presented either on all screens simultaneously (all-screen) or on one screen only (contralateral, central, or ipsilateral relative to recording site). **(b)** Mean responses of all visually responsive neurons to natural image stimuli as a function of screen location across regions. Firing rates were normalized relative to each neuron’s baseline activity (150 ms to 0 ms before stimulus onset). All regions showed significantly larger responses to all-screen vs. partial-field stimuli (linear mixed-effects model with stimulus spatial extent as fixed effect and session and neuron as random effects). Error bars represent 95% confidence intervals of the estimated marginal means. **(c)** Responses (mean ± s.e. across trials) of the same example neurons as in (a) to four visual feature categories: grating spatial frequency, orientation, location and natural image. Responses to spatial frequency were averaged across 6 orientations, responses to orientations were averaged across 4 spatial frequencies, responses to grating locations only contained 90° orientation and 1/15 degrees/cycle spatial frequency. Red shaded boxes indicate when stimulus was shown. **(d)** Cumulative distributions of selectivity indices for visually responsive neurons in each region (blue, striatum; green, GPe; orange, SNr). Selectivity index is defined as the maximum response to any stimulus divided by the mean response to all stimuli, with preferred stimulus identified on odd trials and index computed on even trials. Statistical comparisons used linear mixed-effects models with post-hoc pairwise comparisons (estimated marginal means, Holm-Bonferroni corrected).

In naive mice, neurons in all regions showed larger responses when stimuli appeared on all three screens simultaneously (all-screen), compared to when they appeared on a single screen (**Fig. 2a,b**).

Striatal neurons showed significantly higher stimulus selectivity than GPe and SNr neurons (**Fig. 2c,d**). We computed a feature selectivity index for each neuron and stimulus class by first identifying its preferred stimulus within a set (using only odd trials), then computing the cell’s mean response to this stimulus divided by its overall mean response to all stimuli in the set (using only even trials). This index was calculated separately for four stimulus sets, quantifying selectivity to grating spatial frequency, orientation, location, and to natural image identity, for all neurons which significantly responded to at least one stimulus in the set. For all four stimulus sets, striatal neurons were more selective than GPe and SNr neurons (p<0.002, ANOVA) but GPe and SNr were not significantly different (p>0.38, ANOVA).

These findings show that visual information is present across the basal ganglia of naive mice, but undergoes transformation: while striatum encodes stimulus features, GPe and SNr preferentially encode the presence of a visual stimulus and few details about stimulus identity.

### Training in visuomotor tasks increases visual responses across the basal ganglia

We next asked whether neurons in GPe and SNr might develop more selectivity for visual stimuli in mice that had learned to differentially associate these with actions. To do so, we designed two visuomotor tasks in which a common set of stimuli had different motor associations (**Fig. 3a**). The tasks shared the same basic structure: on each trial, one of three visual stimuli appeared on the right-hand screen with its position yoked to wheel movement^42,43^, but stimuli differed in their action contingencies. In the first task (the “All-Go” task, n = 8 mice), all three stimuli were rewarded for Go responses. In the second task (the “Go/No-Go” task, n = 9 mice), stimuli 1 and 2 were rewarded for Go responses, while stimulus 3 was rewarded if movement was withheld.

**Figure 3:**
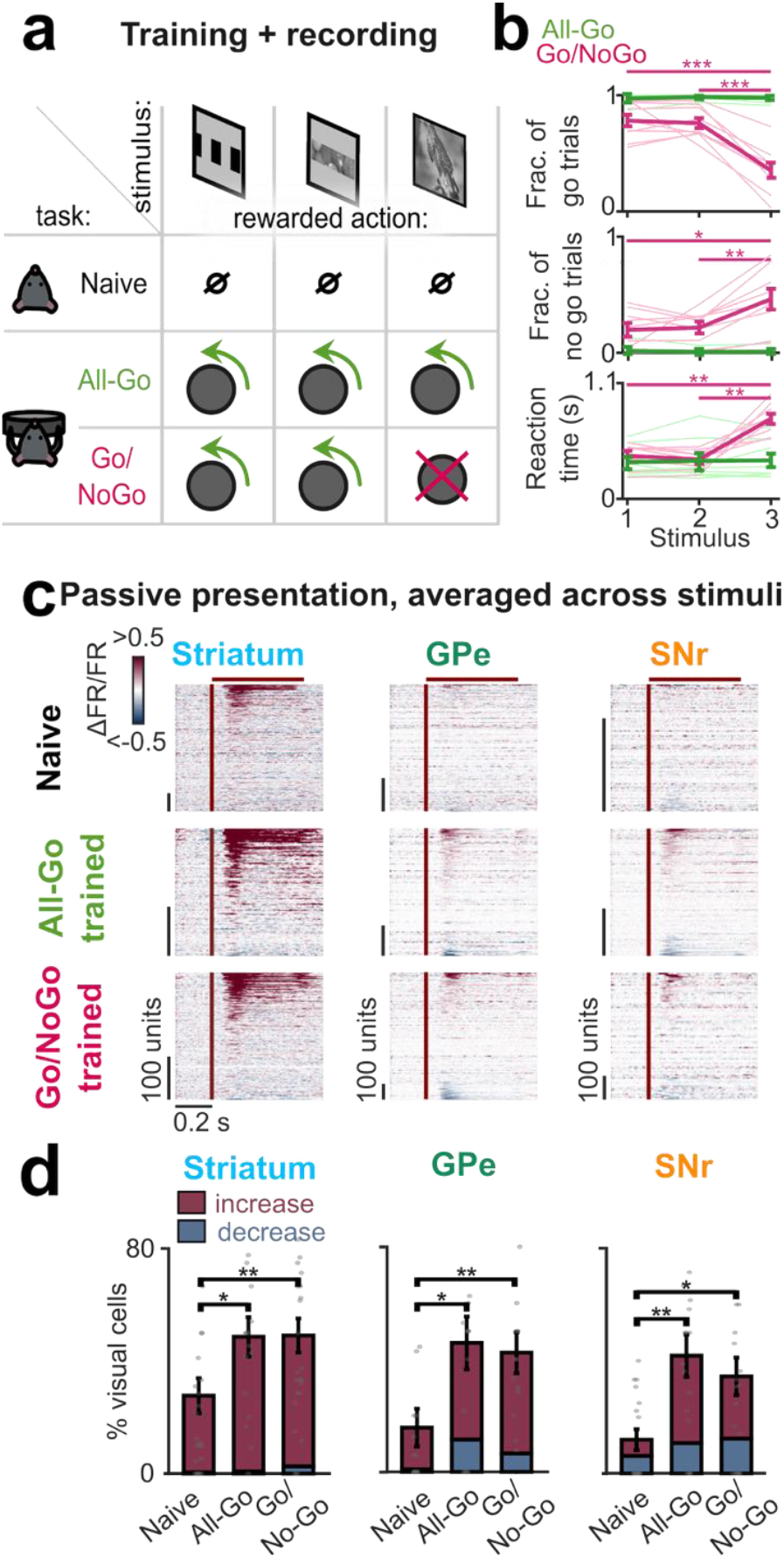
Visuomotor training increases visual responses across the basal ganglia. (a) Schematic of experimental design: mice were either naive (untrained) or trained in one of two tasks: All-Go (green), where all stimuli required a wheel turn response, or Go/No-Go (pink), where two stimuli required turns and one required withholding movement. (b) Behavioral performance. Top panel: probability of “Go” responses to each stimulus in both tasks. Middle panel: probability of “No-Go” responses to each stimulus in both tasks. Bottom panel: reaction times for each stimulus. Green, All-Go task; Pink, Go/No-Go task. Significance stars: paired t-tests, * p < 0.05; ** p < 0.01; *** p <0.001. Error bars: s.e. (c) Pseudocolor raster of neural responses to passive stimulus presentations, for all recorded neurons in each brain region (striatum, GPe, and SNr), in naive mice and mice trained in either task. Red vertical lines indicate stimulus onset. Red horizontal bars indicate stimulus duration. Each cell’s activity is normalized to baseline period mean firing rate (150 ms to 0 ms before stimulus onset). (d) Percentage of cells showing increases or decreases in response to stimulus onset, for each training condition and brain region. Stacked bars show the proportion of cells with increased (red, top) and decreased (blue, bottom) visual responses. Between-task comparisons for total responses (increase + decrease): * p < 0.05, ** p < 0.01. Error bars represent s.e.

Critically, we used identical stimuli across all mice: stimuli 1 and 2 were Go-associated in both tasks, while stimulus 3 was Go-associated in the All-Go task but No-Go-associated in the Go/No-Go task. This design allowed us to examine how behavioral association affects selectivity independent of stimulus identity.

Mice successfully learned both tasks (**Fig. 3b**). In the All-Go task, accuracy exceeded 97% with fast, consistent reaction times for all stimuli. In the Go/No-Go task, mice reliably withheld responses to No-Go stimuli, making Go responses on only 35% of No-Go trials (compared to 77% of Go trials). Incorrect Go responses to the No-Go stimulus had significantly longer reaction times.

Training increased the prevalence of visual responses throughout the basal ganglia (**Fig. 3c-d**). In naive mice, the task stimuli evoked responses in relatively few neurons across all three nuclei, as previously observed for stimuli only appearing on one screen (**Fig. 2b**). After training in any task, however, these same stimuli drove substantially more visual responses across all nuclei. As in naive mice, striatal responses in trained animals were predominantly increases in firing rate, while GPe and SNr showed both increases and decreases (**Fig. 3d**).

The fraction of visually responsive neurons was significantly higher in trained than naive mice across all three basal ganglia regions (**Fig. 3d**). In naive mice, the task stimuli evoked responses in relatively few neurons (28% in striatum, 16% in GPe, 12% in SNr), consistent with the weaker responses to single-screen versus all-screen stimuli observed in naive mice (Fig. 2a,b). After training, these fractions increased substantially: to 49% in striatum, 46% in GPe, and 42% in SNr for All-Go mice, and 49%, 42%, and 34% for Go/No-Go mice. The two task variants did not differ significantly in the fraction of responsive neurons.

### Training increases responses to Go stimuli in all nuclei

Having shown that task training potentiates visual responses across all three nuclei, we next examined whether these enhanced responses encode stimulus identity or behavioral association.

After task training, stimulus representations differed fundamentally across the basal ganglia: only striatal, not SNr or GPe neurons strongly encoded both specific stimulus identities and behavioral association. During passive stimulus presentation after task training, many striatal neurons responded preferentially to specific stimuli even when those stimuli were associated with identical behavioral responses (**Fig. 4a-c**). By contrast, GPe and SNr neurons showed weak stimulus selectivity in the All-Go task, and in the Go/No-Go task they distinguished strongly between Go and No-Go stimuli, but weakly between the two Go stimuli (**Fig. 4d-f**).

**Figure 4:**
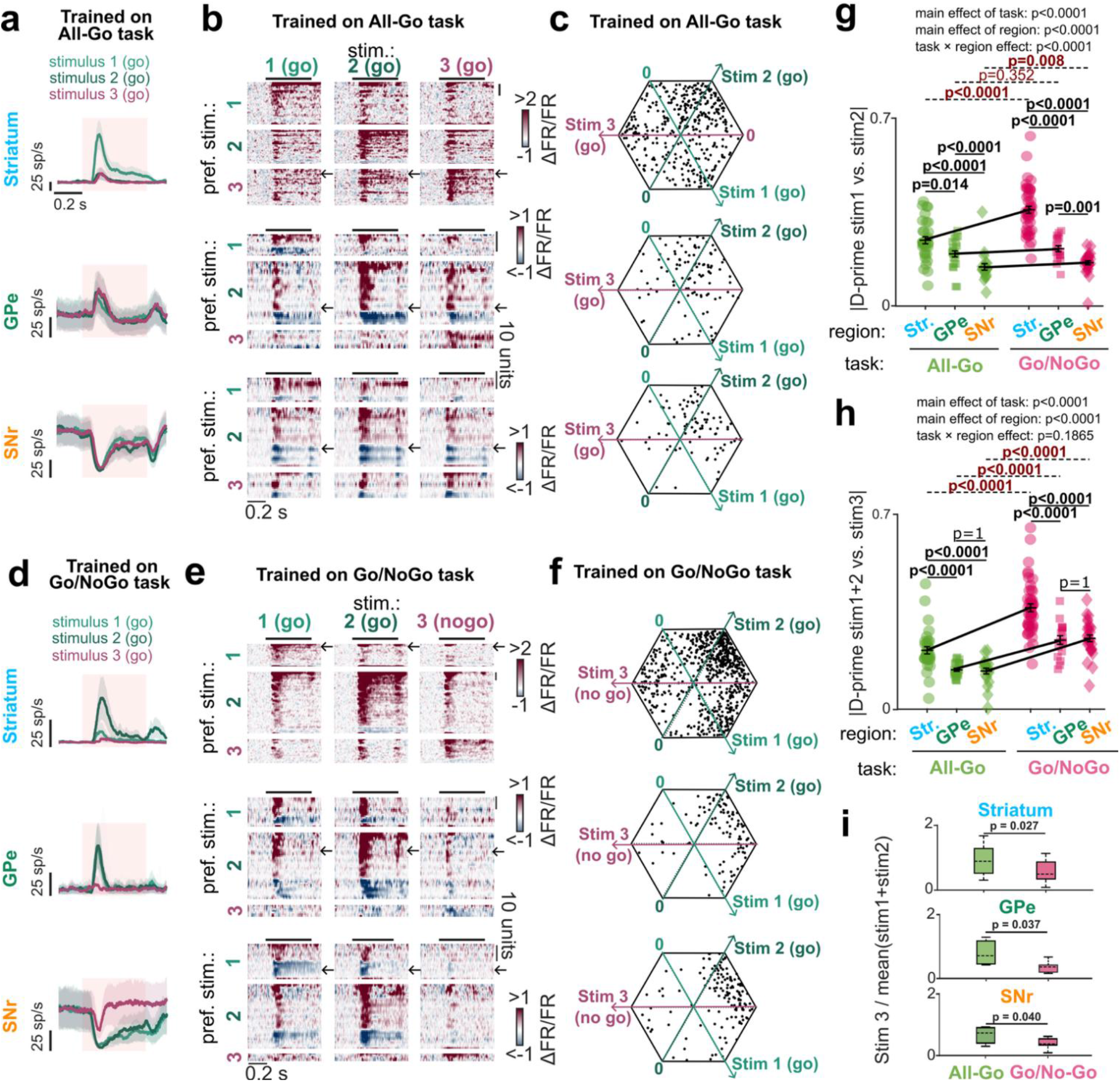
Gradient of stimulus versus behavioral encoding from striatum to GPe/SNr. **(a)** PSTHs showing mean firing rate (solid lines) ± s.e. (shading) of an example neuron from each of striatum, GPe, and SNr in response to passive presentation of the 3 task stimuli, after training on the All-Go task. Colors indicate stimulus identity (light green, stimulus 1; dark green, stimulus 2; dark pink, stimulus 3). Red shaded boxes indicate stimulus duration. **(b)** Pseudocolor raster plot of passive stimulus presentation responses, for all neurons responsive to at least one stimulus from all All-Go-trained mice. Each neuron corresponds to a row; for each brain region, the rows were separated vertically into three bands according to each neuron’s preferred stimulus (i.e. the stimulus producing the largest increase or decrease in rate), resulting in 9 bands altogether (3 preferred stimuli x 3 brain regions). The three columns show these neurons’ responses to the three stimuli, with the same vertical ordering of neurons in each column. Arrows indicate location of example neurons shown in (a). **(c)** Snowflake plots^44^ showing each neuron’s activity to the three stimuli across brain regions for mice trained in the All-Go task. Each point corresponds to one neuron with significant task response. Neurons are plotted in a ternary coordinate system where each vertex represents maximal response to one of the three stimuli (stimuli 1, bottom right; 2, top right; and 3, left), with distance from center indicating response magnitude and angular position indicating stimulus preference. Points closer to vertices show stronger preference for that stimulus, while points near the center show weak or non-selective responses. **(d)** Example neurons from Go/No-Go-trained mice, same format as (a). **(e)** Pseudocolor rasters for Go/No-Go-trained mice, same format as (b). **(f)** Snowflake plots showing each neuron’s activity to the three stimuli across brain regions for mice trained in the Go/No-Go task, same format as (c). **(g)** Discriminability (d-prime) between stimulus 1 and stimulus 2 responses for each session, calculated as d’ = |μ_1_ − μ_2_| / √((σ_1_^2^ + σ_2_^2^) / 2), where μ and σ^2^ are the mean and variance of firing rates in the response window (50–200 ms after stimulus onset) relative to baseline period (−150 to 0 ms before stimulus onset). Each point represents one session. Black lines indicate differences within region across tasks. Error bars represent s.e. p-values: two-way ANOVA. **(h)** Discriminability (d-prime) between Go stimuli and No-Go stimulus for each session. Trials from stimuli 1 and 2 were pooled to compute the mean (μ_Go) and variance (σ_Go^2^) of the Go response. D-prime was calculated as d’ = |μ_Go − μ_3_| / √((σ_Go^2^ + σ_3_^2^) / 2), where μ and σ^2^ are the mean and variance of firing rates in the response window (50–200 ms after stimulus onset) relative to baseline period (−150 to 0 ms before stimulus onset). Each point represents one session. Black lines indicate differences within region across tasks. Error bars represent s.e. p-values: two-way ANOVA. **(i)** Ratio of mean response to stimulus 3 divided by mean response to stimuli 1 and 2, calculated as Ratio = μ_3_ / ((μ_1_ + μ_2_) / 2), where μ_1_, μ_2_, and μ_3_ are mean firing rates in the response window (50–200 ms after stimulus onset). Box plots show median (solid line), mean (dashed line), interquartile range (box), and range (whiskers). In the Go/No-Go task, this ratio was significantly lower than in the All-Go task across all regions. p-values: Mann-Whitney U test.

To test whether striatum encodes visual information even when it is not behaviorally relevant, we computed discriminability indices comparing responses to stimuli 1 and 2, which carried identical behavioral meaning in both tasks (both were always Go stimuli). For each region and session, we computed d-prime for every cell between these two behaviorally identical stimuli and averaged the absolute value across cells (**Fig. 4g**). This index was significantly higher in striatum than GPe or SNr (main effect of region, two-way ANOVA with task and region as factors; p < 0.0001), indicating that striatum more strongly encodes visual differences between stimuli that share the same behavioral consequence. This regional difference was more pronounced in the Go/No-Go than the All-Go task (task × region interaction, p = 0.0001), suggesting that requiring visual discrimination amplifies even task-irrelevant stimulus distinctions specifically in striatum.

Stimuli with different behavioral associations were more discriminable from basal ganglia activity than stimuli with the same behavioral associations (**Fig. 4h**). To show this difference, we recomputed the discriminability index to measure the difference in response to stimulus 3 (whose behavioral significance depended on the task), and the average of stimuli 1 and 2 (which signified Go in both tasks). This information was more strongly encoded in mice that had learned the Go/No-Go task, where it was behaviorally relevant (note the positive slope of black lines in Fig. 4h). Furthermore, the difference between tasks was stronger for this behaviorally relevant information than for the behaviorally irrelevant distinction between stimuli 1 and 2 (compare slopes in Figs. 4h and 4g). To verify this effect statistically, we performed another two-way ANOVA (this time task × selectivity type) for selectivity indices, finding a significant interaction for each basal ganglia region (p < 0.0001). Thus, task training increased stimulus discriminability more when the stimuli were associated with different behavioral requirements, in all basal ganglia regions.

The increased selectivity between Go and No-Go stimuli reflected increased responses to Go stimuli, for all basal ganglia regions (**Fig. 4i**). To measure this increase, we calculated the ratio of the mean response to stimulus 3 (which was associated with either Go or No-Go, depending on the task) divided by the mean response to stimuli 1 and 2 (which were always Go). In the Go/No-Go task, this ratio was significantly lower than in the All-Go task across all three brain regions (p < 0.05 in each region, Ma nn-Whitney U Test), indicating that responses to Go stimuli become stronger than responses to No-Go stimuli, after Go/No-Go training. This specific increase in responses to Go stimuli was even true in the GPe, which would be unexpected under the classical hypothesis that GPe forms a pathway suppressing movement responses.

## Discussion

Our results demonstrate visual responses along all levels of the basal ganglia -- in striatum, globus pallidus external (GPe), and substantia nigra pars reticulata (SNr) -- even in naive mice, but that task training increases these responses, particularly to stimuli associated with “Go” movement responses.

In naive mice, we found visual responses in specific subregions of the striatum, GPe, and SNr that matched an anatomical pathway receiving projections from visual cortices. Visuomotor training increased visual responsiveness across all three regions. Striatal neurons were selective to specific visual stimuli both in naive and in trained mice, and even distinguished stimuli associated with the same behavior. In contrast, neurons in the GPe and SNr were less selective: they responded approximately equally to all stimuli associated with movement, but less to stimuli not associated with movement. Thus, responses in the GPe and SNr were selective to the association of the stimulus, rather than to the sensory features of the stimulus itself. These findings suggest that stimulus information is transformed from stimulus details into behavioral relevance as it flows through the basal ganglia.

### Visual responses in naive mice

Visual responses in the striatum, GPe, and SNr were localized to subregions downstream from visual cortices, matching known anatomical projections from the cortex to striatum and within the basal ganglia^23,25,45–47^. Seminal work in primates established that basal ganglia neurons respond to visual stimuli during saccade tasks^48–55^, but whether such responses exist independent of trained visuomotor associations remained unclear. Our findings demonstrate that visual responses are present throughout the basal ganglia even in naive mice with no task training, suggesting that sensory coding in these structures does not require prior establishment of stimulus-action contingencies. Our findings of visual coding across basal ganglia subregions in the absence of task demands are consistent with an emerging view that basal ganglia structures are not just motor centers^16,56–61^. The visually responsive subregion we identified corresponds to dorsomedial striatum, distinct from the tail of the striatum, which receives input from sensory cortices via a separate anatomical route and has been associated with distinct dopamine signals^62–66^. The functional differences between these visual subregions of the striatum remain to be explored.

What functional role might these visual responses serve in naive mice where stimuli are not associated with behavioral responses? We consider three possibilities. First, natural exploratory behavior might have established weak visuomotor associations^67^, with resulting movements below our detection threshold or occurring outside our measurement window. Second, basal ganglia responses to stimuli in naive mice might represent motor commands which are too weak to effectively drive downstream motor structures, representing a signal that is present but functionally inert. Third, these responses could represent “readiness” for behaviorally relevant associations^68^: baseline sensory representations that do not drive movements themselves, but can be rapidly potentiated once stimulus-action contingencies are established.

Regardless of their specific function, the existence of robust sensory processing in structures traditionally considered motor-specific has clinical implications. Visual deficits often emerge in diseases of the basal ganglia before motor symptoms, including impaired contrast sensitivity and visual discrimination deficits in Parkinson’s disease, and deficits in visual processing and saccadic eye movements in Huntington’s disease^69,70^. Our findings suggest that these early visual symptoms may reflect primary pathology in basal ganglia sensory circuits rather than secondary consequences of motor dysfunction.

### Visual selectivity in naive mice decreases along the basal ganglia nuclei

Stimulus selectivity decreased systematically from striatum to GPe and SNr, matching the anatomical convergence across these structures. In naive and trained mice, striatal neurons showed significantly higher visual feature selectivity than GPe or SNr, paralleling the massive synaptic convergence from striatum to downstream nuclei^23–25^. This transformation from stimulus-specific to non-specific representations is an example of dimensionality reduction, which has been proposed as a core corticostriatal computation^32^. Indeed, if GPe/SNr neurons unselectively sum the activity of many individually selective striatal neurons, GPe/SNr activity would signal that a stimulus is present, but not which stimulus.

### Potentiation of visual responses after visuomotor training

Visuomotor training potentiated visual responses throughout the basal ganglia. Previous work demonstrated that learning potentiates sensory responses in striatum^6–10^. We now show this potentiation extends through the entire circuit to GPe and SNr. The increase in visual responses across the basal ganglia is likely to reflect potentiation of corticostriatal synapses^6^. Increased responses in the GPe and SNr might either be inherited from the striatum or reflect additional plasticity of synapses downstream.

Training mice on a visuomotor association task did not potentiate visual responses uniformly. Go-associated stimuli showed substantially stronger response potentiation than No-Go stimuli across all three nuclei. The basal ganglia may therefore preferentially respond to stimuli associated with actions, consistent with proposals that basal ganglia circuits contribute to the control of movement vigor^71,72^. An alternative possibility is temporally discounted reward expectation: the No-Go stimulus was associated with both lower reward probability (mice were less frequently correct on No-Go trials) and longer delay to reward (1.8 vs. 0.4 s), both of which reduce the temporally discounted reward magnitude of No-Go vs. Go stimuli. Disentangling which of these factors is responsible for the difference between Go and No-Go stimuli would require task designs that orthogonalize reward probability, reward timing, and action requirements.

### Training reshapes the nature of selectivity across nuclei

After training, striatal neurons remained more selective than GPe or SNr neurons, but the form of the difference changed. Striatal neurons in trained mice encoded both stimulus identity and behavioral relevance, distinguishing between stimuli even when they shared identical behavioral associations. In contrast, GPe and SNr neurons responded similarly to all Go-associated stimuli. Thus, the basal ganglia do not simply discard information through convergence; they selectively retain behaviorally relevant features while discarding behaviorally irrelevant features.

This is, to our knowledge, the first direct demonstration that sensory-feature selectivity in striatum is transformed into behavioral-relevance selectivity in downstream basal ganglia nuclei. While previous work has separately documented stimulus-specific responses in striatum^34–37^ and value- or action-related responses in SNr^38,39^, our study directly compared representations across nuclei in the same animals, revealing how information is reorganized along this pathway. This transformation may be fundamental to how the basal ganglia contribute to action selection: by compressing rich sensory representations into signals that reflect what to do rather than what was seen.

We can only speculate as to the mechanism of this specific type of plasticity but hypothesize two possibilities. First, corticostriatal plasticity might strengthen striatal responses specifically to Go stimuli, with this enhanced response conveyed to GPe/SNr via fixed synapses. Second, synaptic plasticity at striatopallidal or striatonigral connections could enhance downstream responses, and if this plasticity were stronger when mice performed an action, it could enhance behavioral distinctions at the expense of stimulus-specific information. The presence of presynaptic D1Rs on striatonigral terminals in SNr^73–75^ raise the untested possibility that dopamine-dependent plasticity at this synapse could exist and contribute to this effect.

### Relationship to direct and indirect pathway function

The view that direct and indirect pathways promote and inhibit behavior remains influential, although it has been modified by recent observations. Activity of direct and indirect pathway neurons is positively correlated: activity in both pathways increases at action initiation^76–78^, and balanced activity across pathways is critical for normal movement^79^. However, when causally manipulated, the pathways exert opponent effects on behavior^80,81^, with the strength of this opponent control depending on task demands and internal state^82^. These results are not contradictory: it is possible that both pathways become active when a subject decides whether to act, with direct and indirect pathways “voting” for and against action^83^, and their co-activation enabling appropriate credit assignment during learning^84^. Our GPe results fit this updated framework, suggesting that GPe activity reflects behavioral relevance rather than uniformly encoding suppression.

This result differs from studies where indirect pathway striatal neurons show enhanced activity during active response suppression^85^ or to unrewarded non-target stimuli^37^. Several factors may account for this discrepancy: the task contingencies differ (active suppression or unrewarded withholding versus our rewarded No-Go condition), we recorded GPe neurons rather than pathway-identified striatal neurons, and our analyses used passive stimulus presentations rather than active task performance. More broadly, our findings add to growing evidence for functional heterogeneity in GPe beyond its classical role in movement suppression^86–88^. Both GPe and SNr contain multiple cell types^22,89–95^; future work could identify which of these is involved in Go-stimulus encoding.

### Summary

In summary, our findings suggest that sensory-driven activity converges across the basal ganglia to turn a stimulus-specific response into a signal more broadly reflecting behavioral relevance. Together, these findings reinforce the fundamental role of the basal ganglia in transforming stimuli into actions.

## Methods

All experiments were conducted according to the UK Animals (Scientific Procedures) Act 1986 under personal and project licences issued by the Home Office.

### Mice

Mice were adult (6 weeks or older) male and female wild type C57BL/6.

### Surgical Procedures

Each mouse underwent 2-3 surgeries: an initial head-plate implantation followed by 1-2 craniotomies for acute electrophysiology recordings.

#### Head-plate Implantation Surgery

Mice were anesthetized with isoflurane and administered subcutaneous carprofen for analgesia before being secured in a stereotaxic apparatus on a heated pad. After shaving the head and disinfecting with iodine and alcohol, the scalp was removed to expose the skull. The skin edges were sealed with VetBond (World Precision Instruments) and the skull surface was thoroughly cleaned. A custom head-plate was attached to the interparietal bone using dental cement (Super-Bond C&B). A 3D-printed polylactic acid U-shaped well was then cemented around the exposed skull perimeter. The skull was protected with a layer of VetBond or Zap-A-Gap (Pacer Technology), followed by two layers of UV-curing optical glue (Norland Optical Adhesives no. 81, Norland Products).

Post-operative analgesia was provided either through carprofen in drinking water for three days or oral Metacam (1.5 mg/ml).

#### Craniotomies

On the first recording day, mice were anesthetized and 1-2 craniotomies (1-2.5 mm diameter) were created using a biopsy punch. The craniotomies were covered with Kwik-Cast (World Precision Instruments) and mice were allowed several hours of recovery before recording began. Between recording sessions, craniotomies were protected with fresh Kwik-Cast applications.

### All-Go task

We designed an operant task where mice used a wheel to move visual stimuli from the right side of a screen to the center to receive water rewards. Water-restricted mice typically received their daily water requirement during task performance or were supplemented to maintain body weight. The task was a variant of one described previously^42^ and programmed using Signals in the Rigbox MATLAB package^96^. Each trial began with one of three visual stimuli (randomly selected) appearing on the right screen: stimulus 1 was a full-contrast grating presented with a spatial frequency of 1/10 degrees/cycle on one-third of the screen (middle portion); stimulus 2 was a natural image of spheres presented in the same location (middle one-third of screen); stimulus 3 was a full-screen natural image of a bird. Natural images were selected from the ImageNet database^97^ from two ethologically relevant categories: ‘birds’ (stimulus 3) and ‘holes’ (stimulus 2). Images were chosen manually to ensure that less than 50% of the image was a uniform background, and to contain a mixture of low and high spatial frequencies. The images were uniformly contrast-normalized. The stimulus position was coupled to wheel movements, with counter-clockwise (leftward) wheel rotations moving the stimulus leftward toward the center. Mice received a 5 μL water reward if they moved the stimulus to center within 1.8 s, after which the stimulus remained centered for 1 s during reward consumption. If mice moved the stimulus rightward off-screen, the trial was aborted and repeated after an extended inter-trial interval (ITI) of 2 additional s. Training progressed through four stages. Stage 1 used a single stimulus with ITIs of 1-3 s and quiescence periods of 0.5-1 s, both selected in 100 ms increments. Stage 2 maintained the single stimulus but extended ITIs to 4-7 s and quiescence periods to 0.5-2 s. Stage 3 introduced the 1.8 s time limit for trial completion. Stage 4 added the second and third stimuli, with all three presented in random order. Training days were not always consecutive, and mice received supplemental water in their home cages on non-training days.

### Go/No-Go task

This task extended the All-Go paradigm by introducing a “No-Go” stimulus requiring response inhibition. Water-restricted mice received their daily water requirement during task performance or were supplemented to maintain body weight. The task was programmed using Signals in the Rigbox MATLAB package^96^. The same three stimuli were used as in the All-Go task: stimulus 1 (full-contrast grating on middle one-third of screen), stimulus 2 (natural image of spheres on middle one-third of screen), and stimulus 3 (full-screen natural image of a bird).

Each trial began with one of three stimuli appearing on the right screen. For the two “ Go” stimuli, the task mechanics were identical to the All-Go task: wheel movements controlled stimulus position, rewards were delivered for successful center placement within 1.8 s, and rightward movements resulted in trial abortion and repetition after an extended ITI. For the “ No-Go” stimulus, the wheel-stimulus coupling remained active, but any movement exceeding 1 degree aborted the trial with an extended ITI and trial repeat. Mice received a 5 μL reward for maintaining the stimulus position within 1 degree for the full 1.8 s. In all rewarded trials, the stimulus remained centered for 1 s during water consumption.

Training progressed through five stages. The first three stages were identical to the All-Go task. Stage 4 introduced the No-Go stimulus, requiring mice to keep the wheel still for 1.8 s to receive a reward. Stage 5 added the second Go stimulus. As with the All-Go task, training occurred on non-consecutive days when necessary, with supplemental water provided on non-training days.

### Passive stimulus presentation

Mice were head-fixed in front of three screens positioned to the left, front, and right, collectively spanning approximately 270 degrees of visual angle in azimuth and 70 degrees in elevation^42,98^.

For naive mice, we habituated them to the rig and head-fixing for at least 4 days, and they were habituated to the full stimulus set for at least 3 days prior to recording. Naive mice were not water-restricted. Recording sessions lasted approximately 2 hours and included four stimulus sets: all-screen static gratings spanning all three screens with varying spatial frequency (1/7.5, 1/10, 1/15 and 1/30 degrees/cycle) and orientation (0°, 30°, 60°, 90°, 120° and 150°); rectangular static gratings at different screen locations; all-screen natural images across all three screens; and a combination of diverse gratings and natural images presented on single screens. Mice could have up to 2 consecutive recording sessions per day. Each stimulus appeared for 500 ms with inter-trial intervals randomly distributed between 0.4-0.6 s. All stimulus sets were repeated at least 50 times, with stimuli presented in a newly randomized order for each repetition.

For trained mice, we presented a subset of visual stimuli passively, consisting of diverse gratings and natural images that included task-relevant stimuli, shown on single screens only. This passive presentation occurred during three acclimation days before training began and after each task performance session once mice were fully trained. Stimuli were presented for 500 ms with inter-trial intervals of 0.4-0.6 s (selected in 100 ms increments) and in randomized order for each session. All stimuli were shown at least 50 times. Trials with any wheel movement were excluded from analysis.

### Neuropixels recordings

We performed recordings using Neuropixels Phase 3A, 1.0, or 2.0 probes^99,100^ mounted on metal rods and positioned with Sensapex micromanipulators. The probe shank angle relative to the base was carefully adjusted for proper insertion. Probes were sharpened using a Narishige EG-45 grinder prior to use to allow for easy penetration of the mouse dura.

To synchronize electrophysiological recordings with task events, we aligned data to a common digital signal that randomly alternated between high and low states, generated by an Arduino. This synchronization method accounted for both clock offset and drift.

Recordings from naive mice occurred during passive visual stimulus presentation only. For trained mice, recordings were conducted during the All-Go task, the Go/No-Go task, and passive stimulus presentations. We inserted a minimum of two probes simultaneously per session. In some cases, we used dual-probe assemblies (two probes cemented together), enabling simultaneous recordings from up to four probes.

### Neuropixels processing

Raw action potential band data were high-pass filtered at 300 Hz. We used CatGT (https://billkarsh.github.io/SpikeGLX; v.3.3) to align channels to each other and apply global common average referencing (-gblcar). Data were then spike-sorted using Kilosort 2 (www.github.com/MouseLand/Kilosort2)^101^. High-quality single units were identified using Bombcell^40^ (https://github.com/Julie-Fabre/bombcell).

Raw waveforms were extracted for each unit (100 spikes per unit) and detrended by removing the best-fit line from each spike. Spatial decay of waveform amplitude across channels was computed and normalized relative to the maximum amplitude. Refractory period violations were calculated using the Wyngaard and Llobet et al. method^102^ with a refractory period of 2 ms and a censored period of 0.1 ms to prevent counting duplicate spikes. Presence ratio was computed using 60 s bins. Percentage of spikes missing was estimated from the amplitude distribution.

Units were classified into three categories: noise, non-somatic, and somatic (good or multi-unit activity). Noise units were identified based on waveform features: maximum of 2 peaks, maximum of 1 trough, waveform duration between 100–1150 μs, maximum baseline fraction of 0.3 (baseline absolute value could not exceed this fraction of the waveform’s absolute peak value), spatial decay slope ≥ −0.008 a.u./μm (linear fit) or between 0.01–0.1 a.u./μm (exponential fit), and second peak-to-trough ratio < 0.8. Non-somatic units^103,104^ were identified by: first peak-to-second peak ratio > 3, main peak-to-trough ratio < 0.8, minimum width of first peak of 4 samples, minimum width of main trough of 5 samples, and trough-to-second peak ratio > 5. Peak detection used a minimum prominence threshold of 0.2 times the maximum waveform value.

Somatic units were further classified as good single units or multi-unit activity (MUA). Good single units met all of the following criteria: minimum amplitude of 40 μV, refractory period violations < 10%, percentage of spikes missing < 20%, minimum of 300 spikes, drift < 100 μm, presence ratio > 0.7, and signal-to-noise ratio > 5. Signal-to-noise ratio was calculated using a baseline noise window of 20 samples. Units failing any of these criteria but passing noise and non-somatic filters were classified as MUA.

### Behavior preprocessing

In naive and trained mice, any trials with wheel movements were excluded from all neural analyses. Wheel movements were detected using a displacement threshold of 0.1 mm. Trials were excluded if the wheel displacement exceeded this threshold for a continuous period of at least 50 ms.

### Histology and electrophysiological alignment

After completing all recordings, mice were deeply anesthetized and transcardially perfused with phosphate-buffered saline followed by 4% paraformaldehyde. Brains were extracted, post-fixed overnight, and embedded in 5% agarose (A9539, Sigma).

Brains were imaged using a custom-built serial-section two-photon microscope. Coronal sections were cut at 50 μm thickness using a vibratome (Leica VT1000), with optical sections acquired every 12.5 μm. Scanning and image acquisition were controlled by ScanImage v5.5 (Vidrio Technologies, USA)^105^ using a custom software wrapper for defining imaging parameters (https://zenodo.org/record/3631609). Brain sections were aligned to the Allen Common Coordinate Framework using brainreg^106^, which was called through our custom software (www.github.com/Julie-Fabre/YAHT). This custom software also enabled manual tracing of probe trajectories and their alignment to the atlas using electrophysiological landmarks recorded during the experiments.

### Allen connectivity

We analyzed viral injection experiments and projection images from the Allen Institute for Brain Science (AIBS) Mouse Connectivity Atlas (http://connectivity.brain-map.org/)^41^ to map corticostriatal and striatofugal pathways. All analysis code has been made freely available at: https://github.com/Julie-Fabre/brain_street_view.

#### Visual Cortex to Striatum projections experiment selection

We queried the API for experiments from wild-type mice (n = 45) and several Cre lines targeting specific cortical layers, focusing on layer 5 and layers 2/3, since these layers project to striatum^107^. The Cre lines analyzed included: A930038C07Rik-Tg1-Cre (layer 5b neurons, n = 41)^108^, Rbp4-Cre_KL100 (layer 5 pyramidal neurons, n = 33)^109^, Tlx3-Cre_PL56 (layer 5a IT neurons, n = 41)^109^, Cux2-IRES-Cre (layers 2/3 and 4, n = 41)^110^, Emx1-IRES-Cre (cortical excitatory neurons, n = 40)^111^. In total, 241 experiments were analyzed across visual cortical areas (VISal: 13, VISam: 21, VISl: 28, VISli: 7, VISp: 150, VISpl: 7, VISpm: 15).

#### Striatum to GPe and SNr projections experiment selection

We analyzed striatal projection patterns to GPe and SNr using experiments with injections in visual-cortex-recipient dorsomedial striatum. All injections were manually inspected before inclusion, excluding any that were mislabeled or contained off-target fluorescence. We manually selected 13 experiments in the GPe and 8 in the SNr including: wild-type mice (n = 2), Drd1a-Cre mice (D1-expressing neurons, n = 3), Plxnd1-Cre mice (n = 3), and Efr3a-Cre mice (n = 1). In addition, for projections from striatum to GPe, we added Drd2-Cre mice (D2-expressing neurons, n = 5). Full experimental details can be found in the Supplementary Table 1.

#### Experiment analysis

For each experiment, we first normalized projection intensity by dividing by the total injection intensity in the source cortical or striatal structure. Specifically, we used data provided by the Allen Institute to normalize: the value ‘sum_pixel_intensity’ given by structure_unionizes for the injection region (https://allensdk.readthedocs.io/en/latest/unionizes.html). We then averaged across experiments and scaled the mean projection intensity from 0 to 1.

#### Visual subregions

To define visual-recipient regions of the striatum from the Allen connectivity experiments, we applied a threshold of 20% normalized projection strength from visual cortical areas. Striatal regions receiving projections above this threshold were classified as visual-recipient subregions and included in subsequent analyses of striatal projection patterns to GPe and SNr. This anatomically defined threshold ensures that, independent of our recordings and experimental conditions, we can fairly compare the percentage of visually responsive neurons across different nuclei and behavioral conditions.

### Peri-stimulus time histograms

Peri-stimulus time histograms (PSTHs) were generated for visually responsive neurons using a cross-validated approach to avoid overfitting. Neurons were sorted based on their peak response time calculated from even trials. For visualization, PSTHs were plotted using the independent odd trials with normalized firing rates calculated as:

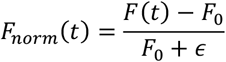

where *F*(*t*) is the firing rate at time t relative to stimulus onset, *F*_0_ is the mean baseline firing rate (-150 to 0 ms pre-stimulus), and ε is a small constant (0.001) to prevent division by zero. This normalization represents the fractional change from baseline while maintaining stability for neurons with low baseline firing rates.

### Classification of visually responsive neurons

To identify and characterize visually responsive neurons, we analyzed neural activity in each region that we identified as visually responsive (see above), aligned to stimulus onset during passive stimulus presentation. All spike data were binned at 1 ms resolution and smoothed with a causal Gaussian filter (σ = 10 ms).

For example cells, we constructed peri-stimulus time histograms showing firing rates aligned to stimulus onset. For population analyses, we created response matrices where neurons were sorted by their mean firing rate during the visual response period (50-200 ms post-stimulus), using cross-validation.

Visually responsive neurons were identified using two criteria: (1) permutation tests comparing firing rates between two time periods: a response period (maximum firing rate 50-200 ms after stimulus onset) and a baseline period (firing rate 150-0 ms before stimulus onset) and (2) *F* = (*r*ˉ[50,200] − *r*ˉ[−150,0])/(*r*ˉ[−150,0] + *ϵ*), where rˉ[t1,t2] denotes the mean firing rate over the time window [t1,t2] ms and with *ϵ* = 0.001. Neurons were classified as visually responsive if their response window activity significantly exceeded baseline activity (p < 0.01, permutation test with 999 shuffles).

For analyses pooling across all stimuli (Fig. 1c–e; Supp. Fig. 4), significance threshold was set at p < 0.01. For analyses requiring visual responsiveness to individual stimuli (Fig. 2b,d; Fig. 3c,d; Fig. 4), the significance threshold was adjusted for the number of stimulus conditions tested (p < 0.01 / number of stimuli) to control for multiple comparisons.

### Naive selectivity index

Stimulus selectivity indices were adapted from Yasuda et al.^38^ to be comparable to their methodology, and calculated for each neuron using a cross-validated approach to avoid overfitting. The dataset was divided into training (odd trials) and testing (even trials) sets.

For each visually responsive neuron (see above) and stimulus s in the training set, we calculated the response strength as:

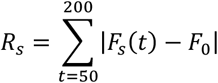

where *F*_*s*_(*t*) is the mean firing rate at time t after the onset of stimulus *s* and *F*_0_ is the corresponding baseline firing rate in the baseline period, 150 ms to 0 ms before stimulus onset. This summed absolute difference captures neurons with temporal coding patterns, such as those exhibiting an initial increase followed by a decrease in firing rate.

The preferred stimulus s* was identified as *s*^∗^ = *argmax*_*s*_ *R*_*s*_ and the selectivity index was then calculated on the independent testing set as:

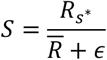

where 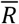 is the mean response to all stimuli and ε = 0.001.

Statistical comparisons of selectivity indices across brain regions were performed using linear mixed-effects models with region as a fixed effect and recording session as a random effect to account for within-session correlations.

#### D-prime

To quantify neural discrimination between task-relevant stimuli in trained mice, we calculated d-prime (d’) for each recording session by averaging across all visually responsive neurons within that session. For discriminability between stimulus 1 and stimulus 2:

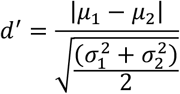

where μ_1_ and μ_2_ are the mean firing rates in the response window (50–200 ms after stimulus onset) for stimuli 1 and 2, and σ_1_^2^ and σ_2_^2^ are the corresponding variances, averaged across neurons within each session.

For discriminability between Go stimuli and the No-Go stimulus:

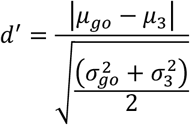

where *μ*_*go*_ and 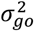 are the mean and variance of firing rates computed across all trials of both Go stimuli (stimuli 1 and 2 pooled together), and μ_3_ and σ_3_^2^ are the mean and variance for stimulus 3. Firing rates were measured in the response window (50–200 ms after stimulus onset) relative to baseline (−150 to 0 ms).

Statistical comparisons of d-prime values were performed using two-way ANOVA with task and brain region as factors. To test whether task training differentially affected behaviorally relevant versus irrelevant selectivity, we performed separate two-way ANOVAs (task × selectivity type) for each brain region, where selectivity type distinguished between the stimulus 1 vs. 2 discriminability index and the Go vs. No-Go discriminability index. Post-hoc pairwise comparisons were performed using Wilcoxon rank-sum tests.

### Stimulus response ratio

To examine how task contingencies affected the relative response to the No-Go stimulus, we calculated for each recording session the ratio of the mean response to stimulus 3 divided by the mean response to stimuli 1 and 2, averaged across all visually responsive neurons within that session:

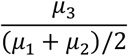

where μ_1_, μ_2_, and μ_3_ are the mean firing rates in the response window (50–200 ms after stimulus onset) for stimuli 1, 2, and 3, respectively.

Statistical comparisons between All-Go and Go/No-Go tasks within each brain region were performed using Mann-Whitney U tests.

## Acknowledgments

We thank Michael Krumin and Flóra Takács for assistance with the experimental setup; Charu Reddy, Laura Funnell, Bex Terry and Yeqing Wang for help with mouse husbandry and training; Magdalena Robacha, George Booth, Isabelle Prankerd, David Orme and Michael Krumin for histology processing; Kevin Miller for helpful discussions and Ilana Witten for reading earlier versions of the manuscript. This work was supported by a Wellcome Trust PhD Studentship, a MathWorks Grant, and by Schmidt Science Fellows, in partnership with the Rhodes Trust, to J.M.J.F, a Wellcome Trust and Royal Society Sir Henry Dale Fellowship (224156/Z/21/Z) to A.J.P., Wellcome Trust Investigator Award (223144/Z/21/Z) to KDH and MC, and ERC Advanced Grant (101097874) to KDH.

## Author contributions

Conceptualization, J. M.J.F., A. J.P., and K.D.H.; investigation, J.M.J.F.; methodology, J.M.J.F.; software, J.M.J.F.; analysis, J.M.J.F.; writing, J.M.J.F., A. J.P., M.C., and K.D.H.; supervision, M.C., A. J.P. and K.D.H.; funding acquisition, J.M.J.F., A. J.P., M.C., and K.D.H.

## Supplementary materials

**Supplementary Figure 1:**
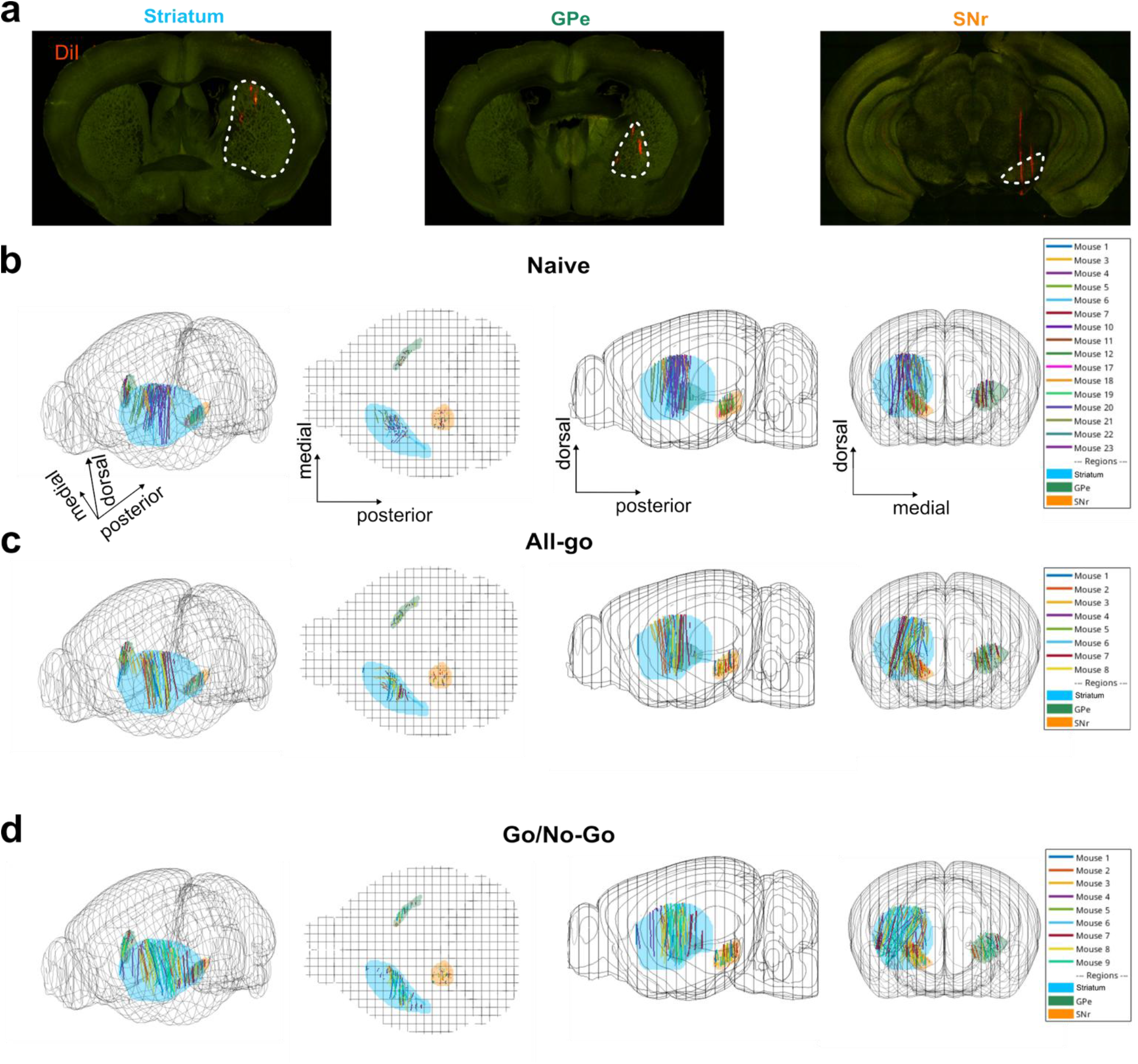
Probe recording locations. (a) Example histology images showing probe tracks (dye-coated probe in red) in striatum (left), GPe (middle) and SNr (right). White outlines indicate each region’s approximate outline. (b) Reconstructed probe tracks for naive mice. Grid represents the brain outline in Allen CCF space, from left: oblique view (azimuth =-37.5°, elevation = 30°), top view, side view, frontal view. Each line represents the trajectory of individual recording probes across multiple mice (n = 23 mice total). Region outlines shown in blue for striatum, green for GPe and orange for SNr. (c) Same as (b) for All-Go – trained mice (n = 8 mice total). (d) Same as (b) for Go/No-Go –trained mice (n = 9 mice total).

**Supplementary Figure 2:**
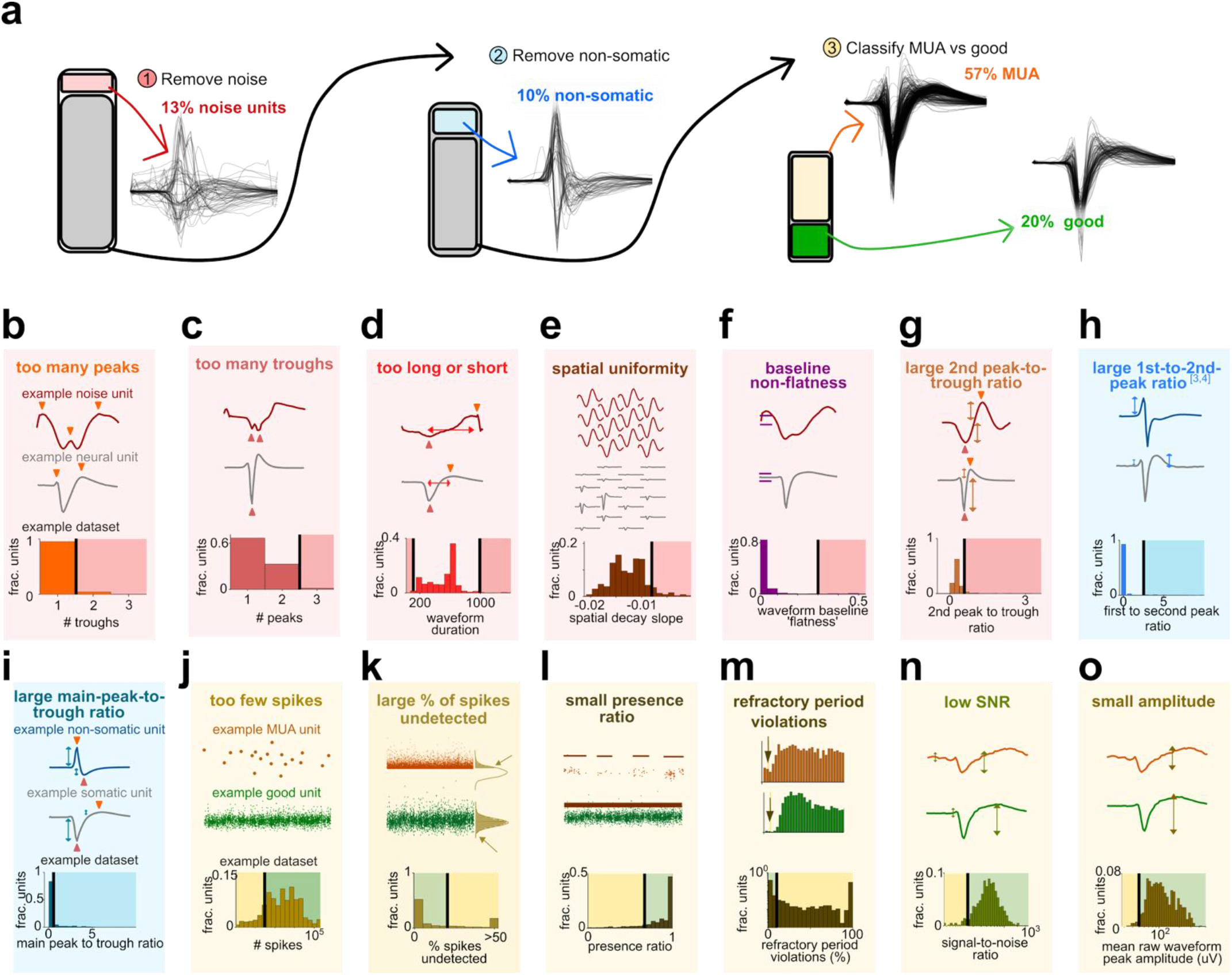
Bombcell quality control pipeline and metric distributions for an example dataset. (a) Schematic of the bombcell automated curation pipeline applied to an example dataset. Raw spike-sorted units undergo three sequential filtering stages: removal of noise units (75 units, 13% of total), removal of non-somatic units (54 units, 10%), and classification of remaining units as multi-unit activity (MUA; 321 units, 57%) or good single units (111 units, 20%). Across all datasets: n = 12,716 total units, of which 2,513 were classified as good single units, 7,604 as MUA, 633 as non-somatic, and 1,964 as noise. (b–o) Distributions of quality metrics used at each filtering stage, shown for the same example dataset. For each metric, top subpanels show example waveforms or unit properties illustrating each classification outcome; bottom subpanels show the distribution of that metric across all units. Units are color-coded throughout by their final classification: green, good single units; gray, somatic/neuronal units not meeting single-unit criteria; orange, MUA; blue, non-somatic; red, noise. Noise detection metrics (b–g): (b) number of peaks in the mean waveform, (c) number of troughs in the mean waveform, (d) waveform duration, (e) spatial uniformity of waveform shape across recording sites, (f) baseline flatness, (g) second peak-to-trough amplitude ratio. Non-somatic detection metrics (h–i): (h) first-to-second peak amplitude ratio, (i) main peak-to-trough amplitude ratio. Single-unit quality metrics (j–o): (j) total spike count, (k) percentage of undetected spikes, (l) presence ratio, (m) refractory period violations, (n) signal-to-noise ratio, (o) mean waveform amplitude.Opus 4.6

**Supplementary Figure 3:**
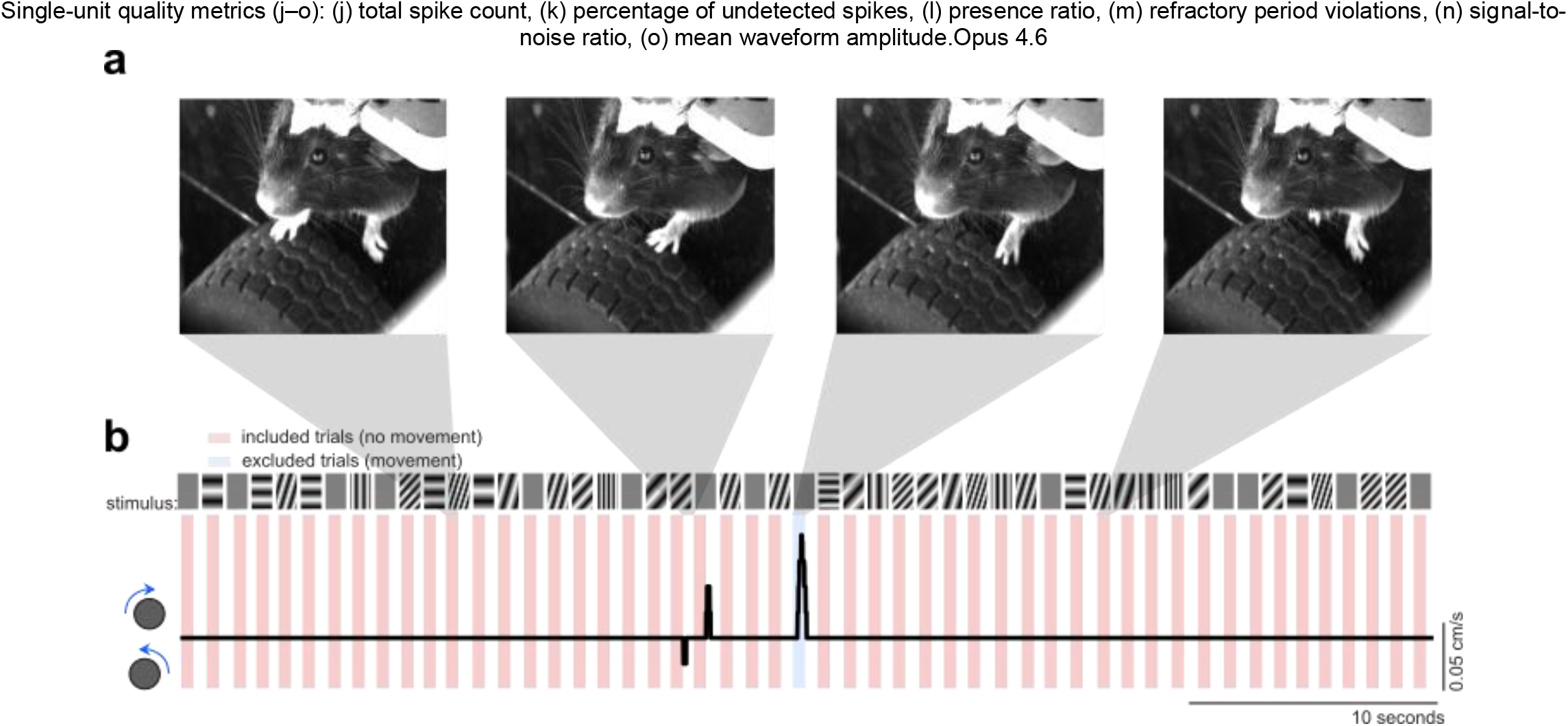
Excluding movements from passive recordings. (a) Sample video frames extracted from behavioral recording at four time points during the session (marked by gray insets in panel b). (b) Example wheel trace (black line) in a naive mouse passively viewing gratings of different spatial frequency and orientation. Timeline of task events and wheel movements from an example session in a naive mouse, showing gratings of various orientation and spatial frequency appearing in front of the mouse (top) and velocity of the steering wheel (bottom). Upward values indicate clockwise wheel movements and downward values indicate counter-clockwise wheel movements. Rectangles indicate stimulus-on times. Included trials with no detectable movement shown in red and excluded trials in blue. Wheel movements were detected using a displacement threshold of 0.1 mm. Trials were excluded if the wheel displacement exceeded this threshold for a continuous period of at least 50 ms. n = 325900 total trials, 13294 excluded trials.

**Supplementary Figure 4:**
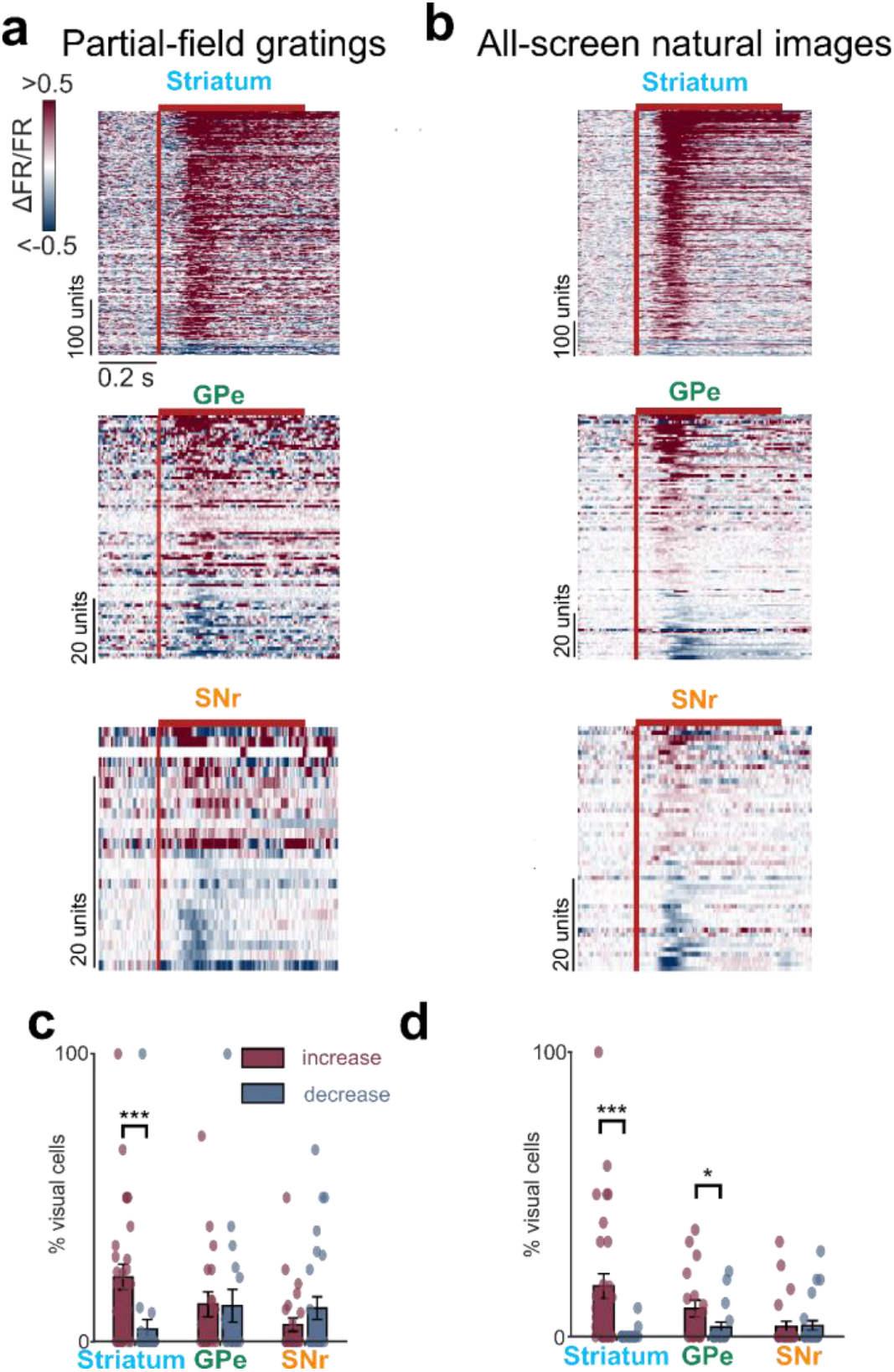
Visual responses by stimulus type. (a) Pseudocolor PSTHs of all visually responsive neurons’ responses to partial-screen gratings of varying spatial frequency and orientations, sorted vertically by response magnitude (average z-scored response 50 to 200 ms after stimulus onset on even trials, odd trials are plotted here). Top: Striatum, middle: GPe, bottom: SNr. Red bar: stimulus duration. Visually responsive neurons were defined with permutation tests between baseline and post-stimulus onset firing rate (p < 0.01, 999 permutations). n = 472 neurons (striatum), 110 neurons (GPe), 51 neurons (SNr). (b) Same as (a) for all-screen natural images. n = 710 neurons (striatum), 110 neurons (GPe), 51 neurons (SNr). (c) Fraction of visually responsive neurons to partial-screen gratings in visual-recipient subregions of each nucleus, separated by response type (increases (red) vs. decreases (blue) in firing rate). Two-way ANOVA (region × response type): main effect of region F(2,140) = 2.270, p = 0.11; main effect of response type F(1,140) = 12.395, p = 0.0006; region × response type interaction F(2,140) = 6.218, p = 0.0026. Post-hoc comparisons for increased responses (Tukey HSD): striatum vs. GPe p = 0.29, striatum vs. SNr p = 0.011, GPe vs. SNr p = 0.49. Within-region comparisons of increases vs. decreases were performed using paired t-tests (Holm-Bonferroni corrected): striatum p = 0.0002, GPe p = 0.39, SNr p = 1. n = 472 neurons, 31 sessions, 8 mice (striatum); n = 110 neurons, 18 sessions, 8 mice (GPe); n = 51 neurons, 24 sessions, 6 mice (SNr). Error bars represent s.e. (d) Same as (c) for all-screen natural images. Two-way ANOVA (region × response type): main effect of region F(2,142) = 2.341, p = 0.10; main effect of response type F(1,142) = 18.217, p < 0.0001; region × response type interaction F(2,142) = 10.750, p < 0.0001. Post-hoc comparisons for increased responses (Tukey HSD): striatum vs. GPe p = 0.20, striatum vs. SNr p = 0.0008, GPe vs. SNr p = 0.20. Post-hoc comparisons for decreased responses (Tukey HSD): striatum vs. GPe p = 0.80, striatum vs. SNr p = 0.025, GPe vs. SNr p = 0.19. Within-region comparisons of increases vs. decreases were performed using paired t-tests (Holm-Bonferroni corrected): striatum p < 0.0001, GPe p = 0.011, SNr p = 0.69. n = 710 neurons, 31 sessions, 8 mice (striatum); n = 110 neurons, 19 sessions, 8 mice (GPe); n = 51 neurons, 24 sessions, 6 mice (SNr). Error bars: s.e.

**Supplementary Figure 5.**
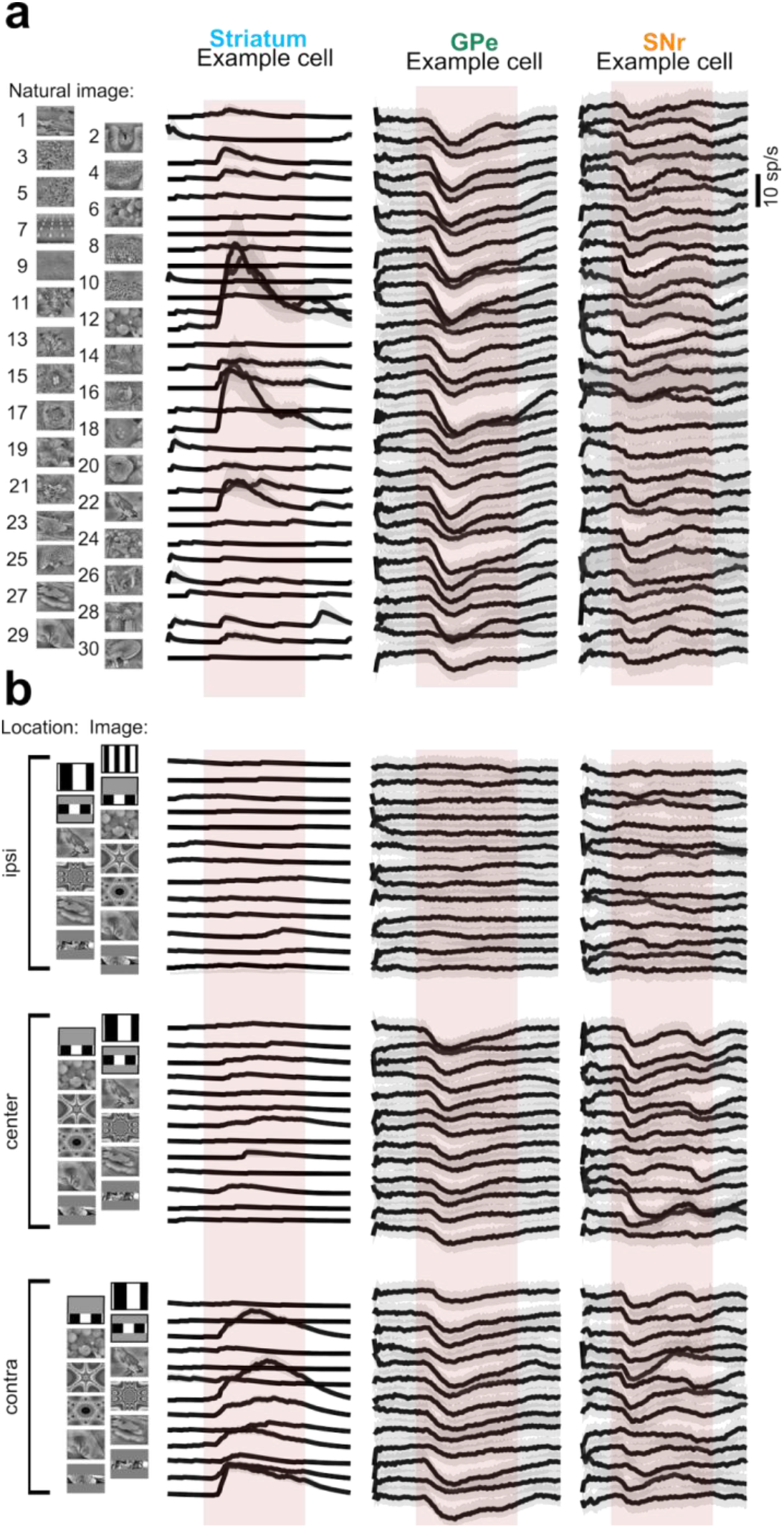
Full stimulus set and example neuron responses. (a) Responses (mean ± s.e. across trials) of the same example neurons as in Fig. 2a,c to natural images. Red shaded boxes indicate when the stimulus was shown on the screen. (b) Same as (a) but for a mix of natural images, fractals, and gratings at different screen locations. Red shaded boxes indicate when the stimulus was shown on the screen.

